# Generation of fluorescent HCV pseudoparticles to study early viral entry events-involvement of Rab1a in HCV entry

**DOI:** 10.1101/2021.03.11.434898

**Authors:** Chayan Bhattacharjee, Aparna Mukhopadhyay

## Abstract

Understanding the early events in viral biology holds the key to the development of preventives. In this study fluorescent Hepatitis C Virus pseudoparticles have been generated where the envelope glycoprotein has a GFP tag. Using these pseudoparticles entry assays were conducted where the entry of the pseudoparticles was tracked via confocal microscopy. Using this system, fusion of host and viral membranes is predicted to occur within 15 minutes of entry in HCV. Using cells with a knockdown for Rab1a, HCV trafficking was observed to be altered, indicating a role of Rab1a in HCV trafficking. In conclusion, this study reports the generation and use of fluorescent pseudoparticles which may be used to understand the early events of viral entry. This system may be adapted for the study of other enveloped viruses as well.

**Highlights:**

- Fluorescent HCV pseudoparticles have been created to study early entry events.
- HCV entry tracking via confocal microscopy reveals fusion within 15 minutes.
- Rab1a is important for HCV trafficking within the cell.

## 1. Introduction

A thorough knowledge of viral entry events hold the key to the development of preventives. Details of viral entry events of enveloped viruses undergoing endocytosis, including Hepatitis C Virus, remain scanty, such as the time it takes internalise and undergo fusion. Hepatitis C virus (HCV) infection is considered to be a global health problem, with more than 70 million infected people worldwide (Lavanchy, 2009) for which so far there aren’t any vaccination. A number of drugs are available although none are preventives and they lack pangenotypic activity coupled with side effects.

HCV, a member of Flaviviridae family encodes for three structural and seven non-structural proteins. The enveloped virus exhibits structural glycoproteins E1 and E2 on its surface of which E2 is a better characterized subunit compared to E1. E1E2 are involved in HCV entry by interacting with various cellular receptors. Following cellular attachment, HCV has been reported to enter hepatocytes via clathrin-mediated endocytosis (Blanchard et al., 2006) which ultimately leads to fusion with endocytic compartments to cause release of the genome into the cytoplasm (Bartosch, Dubuisson, & Cosset, 2003; Hsu et al., 2003). In this endocytosis process, early endocytic sorting and vesicle fission/fusion results in the generation of either recycling endosomes destined for the cell surface or late endosomes that ultimately fuse with lysosomes for degradation (Willingham and Pastan, 1982). Regulation of such sorting events is not completely understood but is believed to be mediated by the small Rab (Ras-related in brain) GTPases (Bananis, Murray, Stockert, Satir, & Wolkoff, 2000; Murray, Sarkar, & Wolkoff, 2008). One such mediator is Rab 1a which is involved in early endocytic sorting for many ligands (Mukhopadhyay et al., 2014) such as ASGPR and EGFR which have been known to be involved in HCV entry (Saunier et al., 2003; Diao et al., 2012).

Details of HCV entry remain elusive and so does an understanding of the timeline of viral entry and membrane fusion event. To address this issue, this manuscript describes creation of fluorescent pseudoparticles for the study of early events of HCV entry. Most of the HCVpps so far used to study HCV entry from different laboratories are GFP/ luciferase reporter based lentiviral particles that have been analyzed via microscopic/ luminescence experiments 72hours of post infection on liver cell lines (Bartosch & Cosset, 2009; Voisset et al., 2006; Baldick et al., 2010). To the best of our knowledge, we report here for the first time fluorescent HCVpp that have been used to study the initial 30 minutes of trafficking events of this virus by fluorescence microscopy.

## 2. Materials and Methods

### 2.1 Cell culture

All experiments were conducted in either HEK 293T, Huh7 or Rab1a KD Huh7 cells. The Rab1aKD cells were obtained as a kind gift from Dr. Allan Wolkoff, Albert Einstein College of Medicine, Bronx, NY. HEK 293T cells were grown in DMEM media and Huh7 cells in RPMI media, containing L-glutamine (Gibco, Thermofisher Scientific, USA), supplemented with 10% FBS and 1% penicillin streptomycin. Rab1a KD cells were maintained similar to Huh7 with an additional supplement of 2.0 μg/ml Puromycin.

### 2.2. Plasmid constructs

Plasmid pcDNA-E1E2 containing the full length cDNA of genotype 1a (H77) strain inserted into pcDNA and retroviral structural genes Gag-Pol containing construct pTG5349, were obtained as a kind gift from Prof. Jean Dubuisson, Université Ede Lille, France. PGK-GFP reporter construct, pCCLSIN.cPPT.hPGK.GFP.WPRE was obtained from Albert Einstein College of Medicine, New York (“HIV-based vectors. Preparation and use - PubMed,” n.d.). EGFP tagged construct was generated in pEGFP-N1 (Clontech).

### 2.3. Antibodies, Fluorescent dyes and probes

Rabbit anti-GFP antibody (G1544) was purchased from Sigma Aldrich, USA (G1544). Plasma membrane marker FM 4-64 lipophilic dye (Molecular probes, Invitrogen detection technologies) and lyso-sensor molecular probe Lyso Tracker (Molecular Probes, Life Technologies) ware purchased from Thermo Fisher Scientific, USA (Thermo Fisher Scientific, USA).

### 2.4. Preparation of fluorescence labelled HCV-E1E2

HCV glycoprotein E1E2 from pcDNA-E1E2 was inserted into pEGFP to prepare a construct expressing E1E2-EGFP fusion protein. For this, E1E2 sequence was amplified via PCR from pcDNA E1E2 vector using primers (forward 5’ CGAAGCTTGCATGGGTTGCTCTTTC 3’. and reverse 5’ CAGAATTCCCGCCTCCGC 3’) the product was subsequently digested with *Hind*III and *Eco*RI (NEB, USA) and ligated into pEGFP-N1 to create a E1E2-EGFP fusion construct with EGFP at the C-terminal end.

### 2.5. Transfection of pEGFPN1-E1E2 in 293T cells for cell surface expression of EGFP

HEK 293T cells were seeded on sterile coverslips to achieve 70-80% confluency. pEGFP-E1E2 vector was transfected using TurboFect (Thermo Fisher Scientific) reagent following manufacturer’s instructions. 48h later, cells were stained with 5μg/ml of plasma membrane marker FM 4-64 for 15min on ice followed by fixation with 4% formaldehyde solution for 10min on ice followed by 10min at room temperature. Overexpression of E1E2-EGFP was checked by confocal microscopy (Leica DMi8) using 63X oil immersion objective. *z*-Series confocal images were acquired with FITC and rhodamine filter settings. All the Microscopic images were quantified by Image J (1.52a) (ImageJ-win-64). Images were pseudocolored and co-localization measured using ImageJ-Fiji image processing software.

### 2.6. Immunoblotting to confirm expression of E1E2-EGFP

To confirm protein expression of pEGFP-E1E2 in HEK293T cells, western blot analysis was performed. Approximately 60% confluent HEK293T cells were seeded on 60×15mm dish (Thermo Fisher Scientific, Denmark) and pEGFP-E1E2 vector was transfected with TurboFect reagent as stated earlier. 48h post transfection cells were scraped and lysed in 100μl lysis buffer containing 50mM Tris, 150mM NaCl, and 0.1% NP40 supplemented with 100mM PMSF (Themo Fisher Scientific, USA), protease inhibitor cocktail (Sigma-Aldrich, USA), protease inhibitor cocktail tablets (Roche, Germany). Lysed cells were centrifuged at 16,000xg for 20min and supernatant stored at −20°C until use. 30μg protein was used per well in 12% SDS-PAGE, followed by immunoblotting with rabbit anti GFP antibody.

### 2.7. Production of fluorescent HCV pseudoparticles (HCVpp)

HCV pseudoparticles were produced by transfecting constructs encoding HCV E1E2-EGFP (E1E2 fused to EGFP), retroviral packaging construct pTG5349 and a reporter pCCLSIN.cPPT.hPGK.GFP.WPRE into HEK 293T cells by calcium-phosphate method (Calphos Mammalian Transfection kit, Clontech as per manufacturer’s instructions). Cells were seeded on 60×15mm dish a day before transfection to achieve 70 – 80% confluency. pTG5349/gag-pol (4.2μg), pCCLsin.cPPT.hPGK.GFP.Wpre (4.2μg) and pE1E2-EGFP (1.4μg) plasmid DNA were mixed with solution A (32μl calcium solution and sterile water up to total volume of 260μl). 260μl of solution B (260μl 2X HBS buffer) was then added to solution A mixture and vortexed slowly for few seconds and incubated at room temperature for 15min. The transfection mixture was added dropwise to HEK 239T cells and allowed to incubate at 37°C in 5% CO2. 3ml fresh DMEM complete media (with FBS) was added after 16h post transfection. After 40h post transfection, HCVpp containing media (approximately 42 ml) was collected, purified via passage through 0.45μm filter (Millipore, MILLEX-GV), concentrated by Amicon ultra filter (Amicon Ultra Centrifugal Filters, Merck) by centrifugation at 4000xg in a swinging-bucket rotor for 20min. The fluorescent HCVpp were stored at −80°C. Approximately 900μl HCVpp is obtained from 42ml of media.

### 2.8 Transduction of Huh7 cells with HCVpp

Confluent Huh7 cells monolayers grown in 100×10mm dish (nuncTM, Thermo Fisher Scientific) were transduced with 90μl HCVpp supplemented with DMEM (910 μl DMEM and 90 μl HCVpp) without FBS and antibiotics on ice for 1h. Infected dish was incubated for 20 min in 5% CO2 at 37°C followed by wash with PBS twice. Post internalization cells were scraped and resuspended in 1ml PBS on ice, followed by a centrifugation at 500xg for 2min. Total RNA was isolated using Trizol reagent (Sisco research laboratories, India) with 400μg Ribonucleoside vanadyl complexes (VRC) from NEB, USA. cDNA was synthesized with 1μg of RNA, using iScript^™^ Reverse Transcription supermix for RT-qPCR kit (BIO-RAD Laboratories, USA) following manufacturer’s instructions at PCR conditions (priming at 25°C for 5min, reverse transcription at 46°C for 20min, inactivation at 95°C for 1min). 60ng of cDNA was amplified by PCR using (20 picomole/ reaction) forward and reverse primer for GFP (forward 5’TCGACAGTCAGCCGCATCT3’ and reverse primer 5’CCGTTGACTCCGACCTTCA3’) respectively using 1μl taq polymerase (NEB, USA). The PCR amplification was performed as per the following cycle: 95°C for 5min, 30 cycles of 95°C for 30sec, 47°C for 45sec and 72°C for 1min followed by a final extension at 72°C for 5 min. The PCR amplified product was checked in 1.5% agarose gel electrophoresis.

### 2.9. Trafficking of EGFP labelled HCVpp in cells

To understand HCV trafficking events from 0-30min, approximately 60-70% confluent Huh7 or Rab1a cells were seeded on sterile coverslips placed in a 60×15mm cell culture dish. 50μl HCVpp diluted in 450μl of DMEM (without FBS, antibiotics) were added (100μl mixture each) dropwise on the coverslips placed on ice. After washes with cold Hanks’ balanced salt solution without magnesium or calcium (HBSS), HCVpp was allowed to internalize by shifting cells to 37°C for various times as required in various experiments. Internalization was stopped by shifting the cells to ice, followed by fixation in 4% formaldehyde for 10min on ice and 10min at room temperature (RT). For staining with a plasma membrane marker FM 4-64, 5μg/ml was incubated for 15min on ice prior to fixation. To study lysosomal colocalisation, Lyso Tracker was added at a concentration of 0.14μM (diluted in PBS) to both of the cell lines for 30min at RT prior to shifting cells to ice and incubation with HCVpp. Cells were examined via confocal microscope (Leica DMi8) in 63X oil immersion and images were quantified. *z-*Series images were acquired confocal microscope utilizing a multichannel white light source, FITC and rhodamine filter settings. Images were pseudocolored and colocalisation measured using ImageJ-Fiji image processing software. Pixel intensities of EGFP or integrated density (IntDen) from all the microscopic images were quantified by Image J (1.52a) (ImageJ-win-64), background IntDen of each of the image was subtracted. All the IntDen of each of the image for all the time points were normalized with mean IntDen of 0min

### 2.11. Statistical analysis

Multiple comparisons were conducted by one-way analysis of variance followed by Bonferonni post hoc analysis. Correlation analysis was performed by linear correlation analysis Pearson’s R (using above threshold values) by Fiji image processing software (ImageJ-win-64). t-score was determined to check the significance of individual Pearson’s R values (above threshold). All statistical analyses were performed using SPSS software (version 17.0; SPSS Inc, Chicago, IL). Difference was considered statistically significant at P < 0.05.

## 3. Results

### 3.1. Preparation of pE1E2-EGFP construct and its expression in 293T cells

E1E2-EGFP fusion protein was conceptualized in order to generate fluorescent pseudoparticles suitable for microscopy studies. A C-terminal EGFP tagged fusion of E1E2 was created such that ER-signal sequence of the protein remained undisturbed. Furthermore as the C-terminal end of E2 is on the cytoplasmic side, an EGFP tag is expected to not significantly affect its ability to interact with the HCV entry receptors.

To check whether the E1E2-EGFP construct was successfully prepared, protein expression of EGFP was analyzed after its transfection into HEK 293T cells. 48h of post transfection EGFP expression was analyzed by microscopic analysis as well as by immunoblotting. EGFP expression was observed to be in vesicular structures throughout the cytoplasm (Fig. 2A). Quantitation of the pixel intensity (IntDen) revealed significantly higher fluorescence intensity signal compared to mock transfected cells (Fig. 2B). EGFP expression was also observed after 48h of post transfection by immunobloting analysis (Fig 2C).

**Fig. 1.**
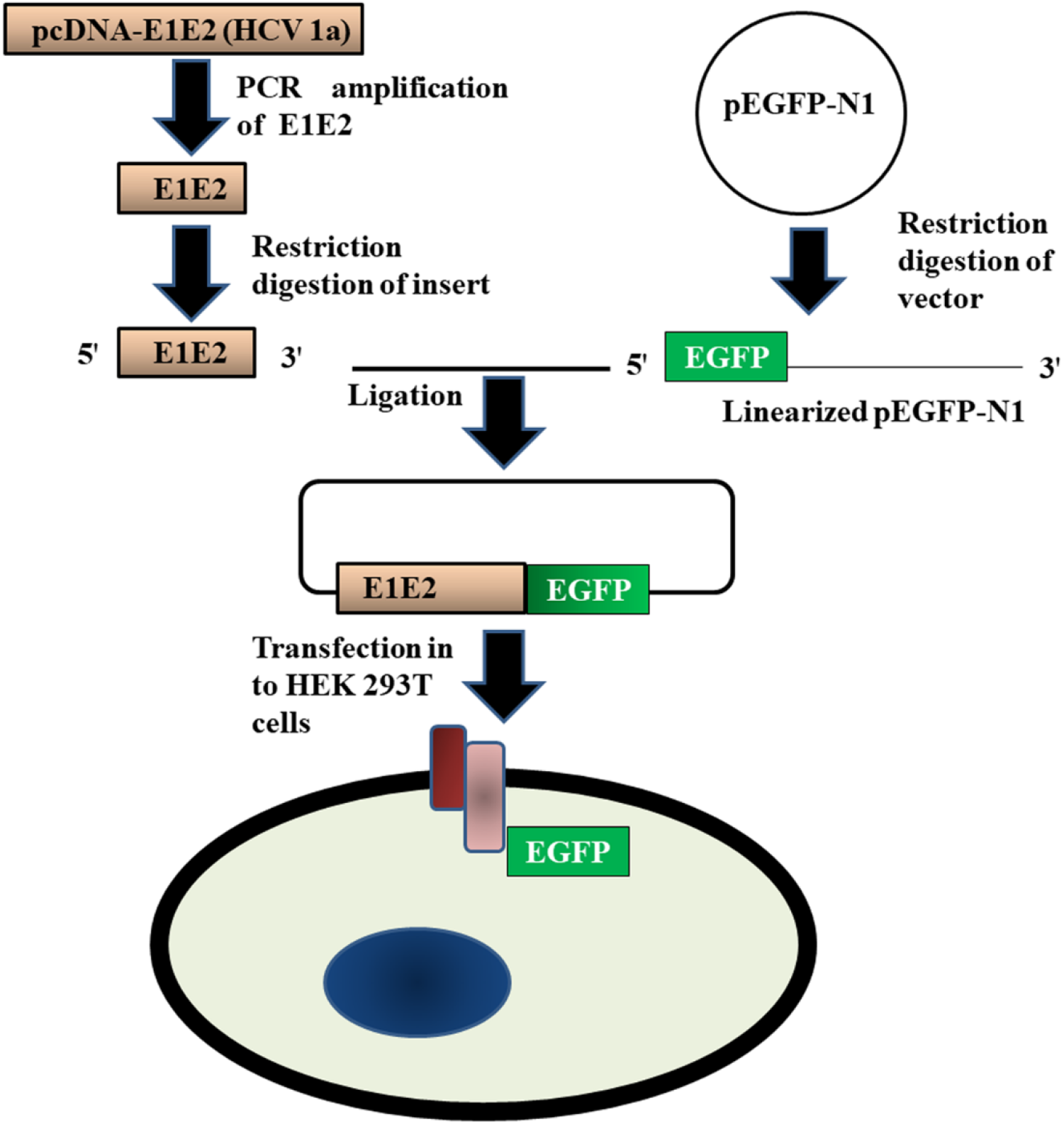
Strategy of generation of E1E2-EGFP fusion protein. E1E2 was PCR amplified, digested with restriction enzymes (EcoR1 and HindIII) and ligated into pEGFP-N1. Transfection of pE1E2-EGFP in HEK 293T cells was expected to yield a fluorescent protein localized on membranes.

**Fig 2.**
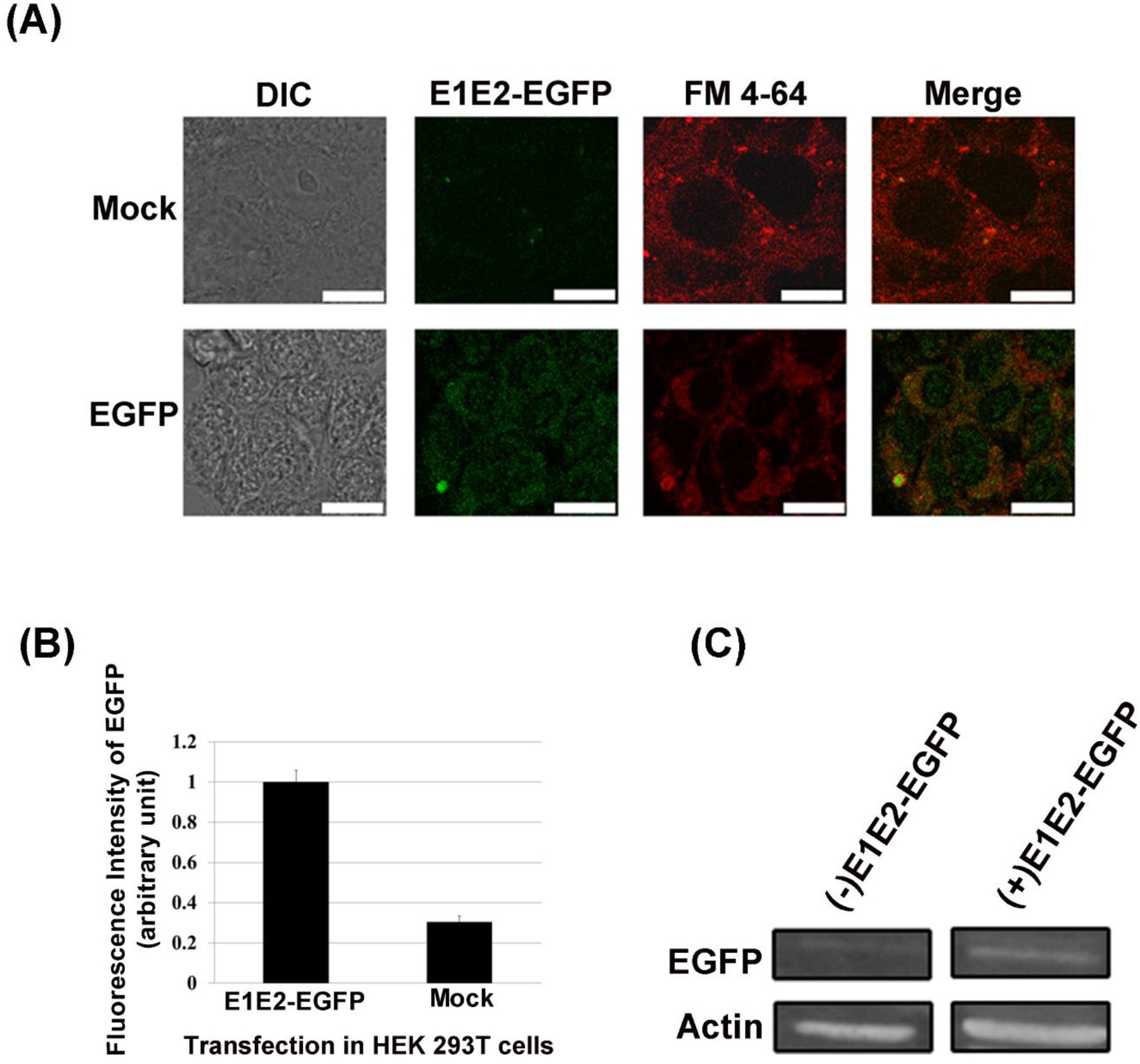
E1E2-GFP fusion construct expression in cells. (A) Representative z-plane confocal images of fixed HEK 293T cells where expression of EGFP (green) in HEK 293T cells after 48h of transfection with pE1E1-EGFP fusion construct was observed. Cells were co-stained with FM4-64 and the merged images depicted. Scale bars= 10μm. (B) Graphical representation of fluorescence intensities of EGFP expression in transfected (E1 E2-EGFP, n =124) and Mock (n=68) in 293T cells (C) Immunoblot for GFP and Actin (control) indicating expression of EGFP in E1E2-EGFP transfected cell lysate.

### 3.2. Preparation of fluorescent pseudoparticles

Lentiviral pseudoparticles (HCVpp) were prepared as depicted schematically in Fig 3 and detailed in the methods section. Briefly, transfection of the E1E1-EGFP, a Gag/Pol packaging construct and a PGK-GFP (pCCLSIN.cPPT.hPGK.GFP.WPRE) containing reporter construct into 293T cells yielded fluorescent pseudoparticles from the media. To check for successful production of HCVP_pp_ in HEK 293T cells, harvested HCVpp were allowed to internalize for 20min in Huh7 cells. EGFP expression was observed under confocal microscope (Fig 4A). Fluorescence was observed in punctate structures throughout the cell. Transduction efficiency was approximately 70%. Presence of PGK-GFP RNA was detected by qRT-PCR from isolated total RNA of the transduced cells which indicated successful production and transduction of pseudoparticles (Fig 4B).

**Fig 3.**
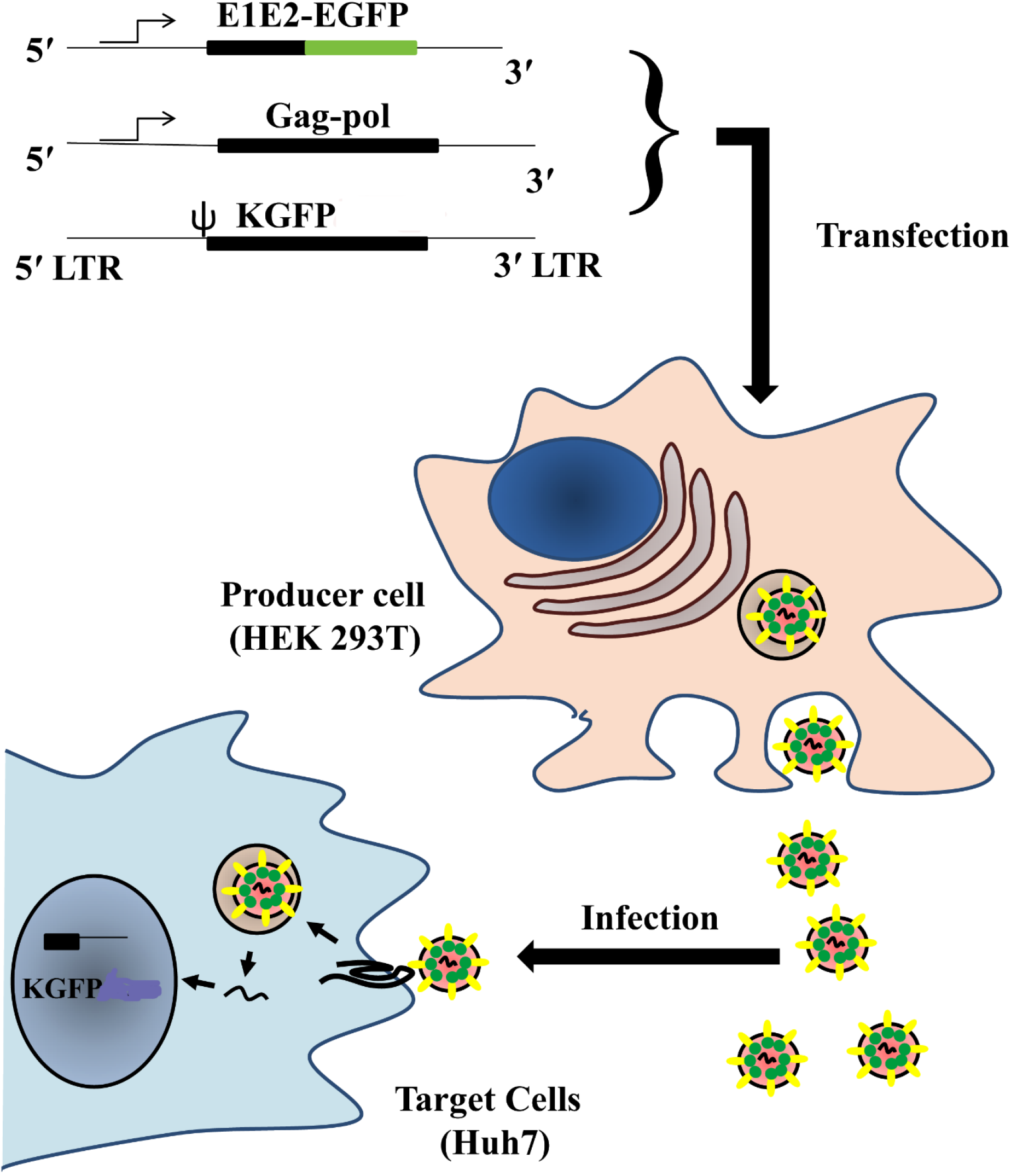
Production of GFP-expressing HCV pseudoparticles (HCVpp). Human embryonic kidney cells (HEK 293T) were transfected with three expression vectors; E1E2-GFP, Gag/Pol packaging construct and pCCLsin.cPPT.hPGK.GFP.Wpre, GFP reporter construct. Fluorescent pseudoparticles extracted from the media of 293T cells were used for infection / entry studies in Huh7 cells and visualised via confocal microscope.

**Fig. 4.**
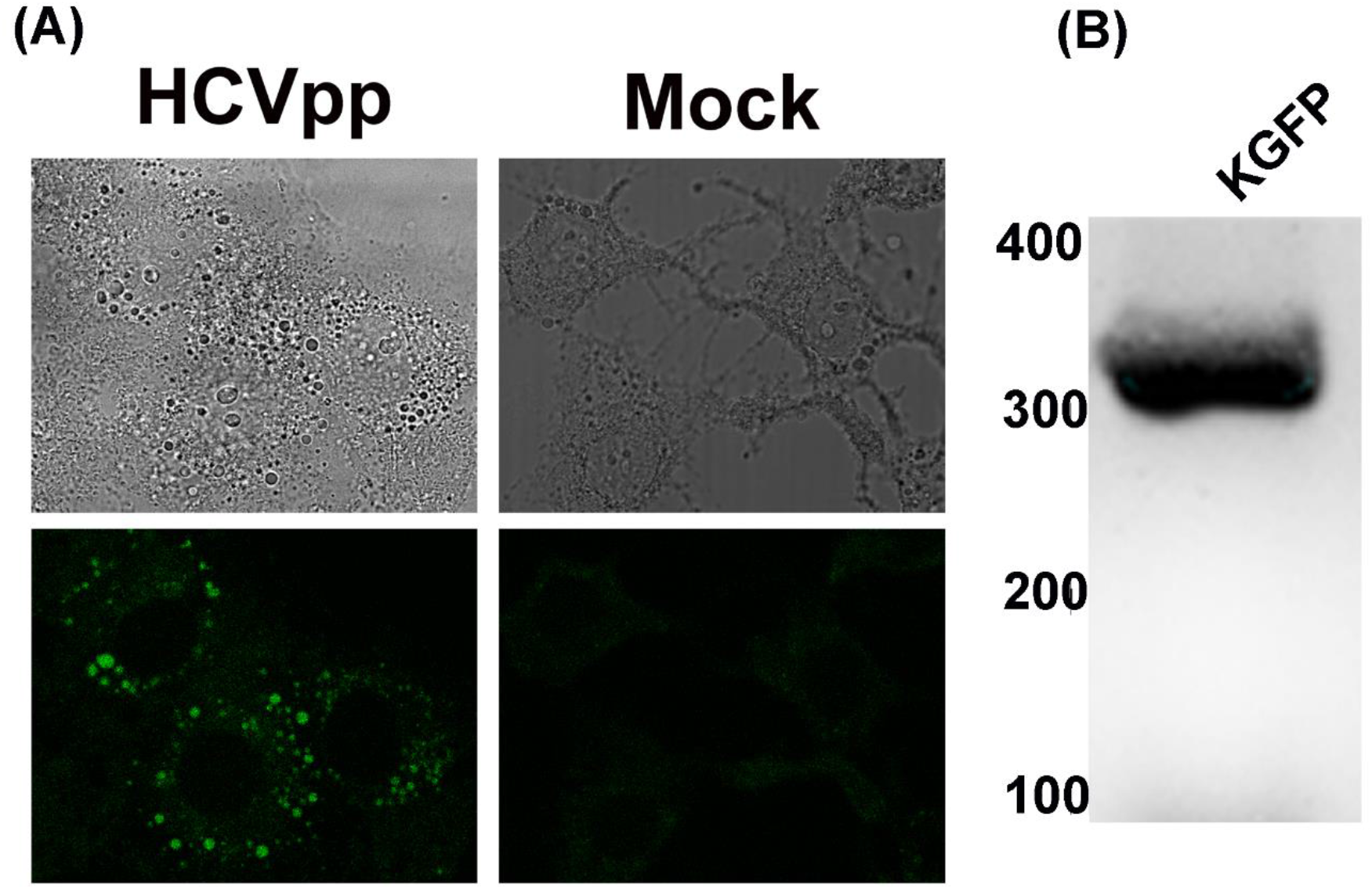
Entry assay for HCVpp in Huh7 cell line. (A) Expression of EGFP in HCVpp infected Huh7 cells via confocal microscopy. (B) RT-PCR amplification of GFP from RNA obtained from Huh7 transduced with HCVpp, separated in 1.5% agarose gel.

### 3.3. HCVpp Trafficking in Huh7

To study HCVpp trafficking, we conducted a time lapse experiment of HCVpp entry in Huh7 cells. As previously characterized, usual endocytic processing in Huh7 results in delivery of ligands to lysosomes within 30 minutes. Hence we developed entry assays to track the entry of fluorescent HCVpp in the initial 30 minutes itself. Fig 5 shows representative images of Huh7 cells that were fixed 0, 5, 10, 15 and 30min after HCVpp internalization.

**Fig. 5.**
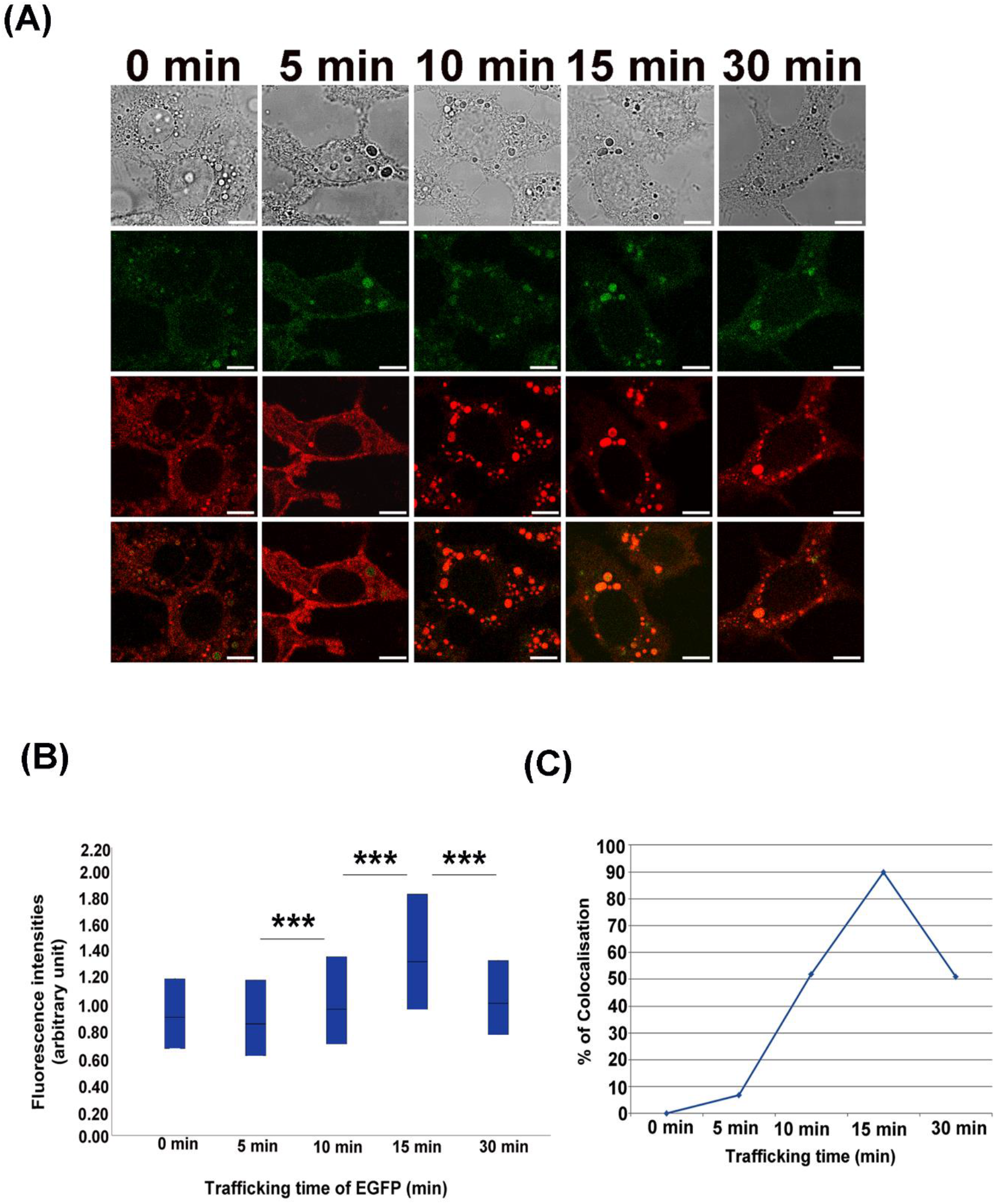
HCV pseudoparticles infection and trafficking in Huh7 cells. Representative z-plane confocal images of fixed Huh7 cells where, (A) Images of HCVpp transduced in Huh7 cells, fixed at indicated times of post internalisation and analysed via confocal microscope. Scale bars= 10 μm. (B) Integrated density (IntDen) of EGFP from each of the individual infected Huh7 cells are represented in a box plot profile. Each box represents the median, first and third quartile. ** represents significant difference (p < 0.05) and *** represents significant difference (p < 0.001) as tested by ANOVA followed by Bonferonni post hoc analysis. For Huh7 (0 min), N= 1748; for 5min, N= 1018; for 10min, N= 768; for 15min, N= 556; for 30min, N= 906. (C) Graphical representation of positive colocalization percentage (%) between EGFP and FM 4-64 observed at indicated time points. N (0 min)= 67, N (5 min)= 46, N(10 min)= 33, N(15 min)= 60 and N(30min)= 76. Total number of IntDen measurements for images throughout a z-stack for all replicates of a time point were counted as N. t-score was determined to check the significance of Pearson’s R (above threshold) values.

Measurement of pixel intensities of EGFP tagged HCVpp in Huh7 is depicted in Fig 5A. It was observed that while pixel intensity remained constant between 0-5 minutes, it rose to a peak level by about 15 minutes. This is possibly due to coalescing of vesicles due to endocytic processing and fusion-fission events. Thereafter the pixel intensity dropped between 15-30 minutes of internalization possibly due to degradation or dissipation of E1E2-EGFP.

To further study the trafficking details, the cells were stained with a plasma membrane dye FM 4-64. Figure 5 shows representative individual and merged images which indicate colocalization between HCVpp and FM 4-64 at various time points. As seen in the images and quantitative data of colocalization between HCVpp and FM 4-64, signals rose sharply to a peak value at 15 minutes. 15 minutes was also the time when maximum pixel intensity of HCVpp was being observed (Fig. 5B). As FM 4-64 is a plasma membrane dye that is not internalized, a high colocalization at 15 min is indicative of trafficking of E1E2-EGFP to the plasma membrane. This would indicate that viral and host membrane fusion has been completed by 15min of internalization and hence the E1E1-EGFP has trafficking back to the cell surface via recycling endosomes. However by 30 minutes a drop in colocalization as well as E1E2-EGFP pixel intensity is perhaps indicative of either dissipation of E1E2-EGFP or its degradation. To cross check degradation in lysosomes, we used a lysosome marker Lyso Tracker and conducted similar type of time lapse experiment. Interestingly we did not find any colocalization even at 30min of internalization in Huh7 (Fig 7.). This finding indicates that E1E2 is not degraded in lysosome after 30 min of internalization. Hence it is possible that the E1E2 dissipated or was degraded via some other mechanism.

**Fig. 6.**
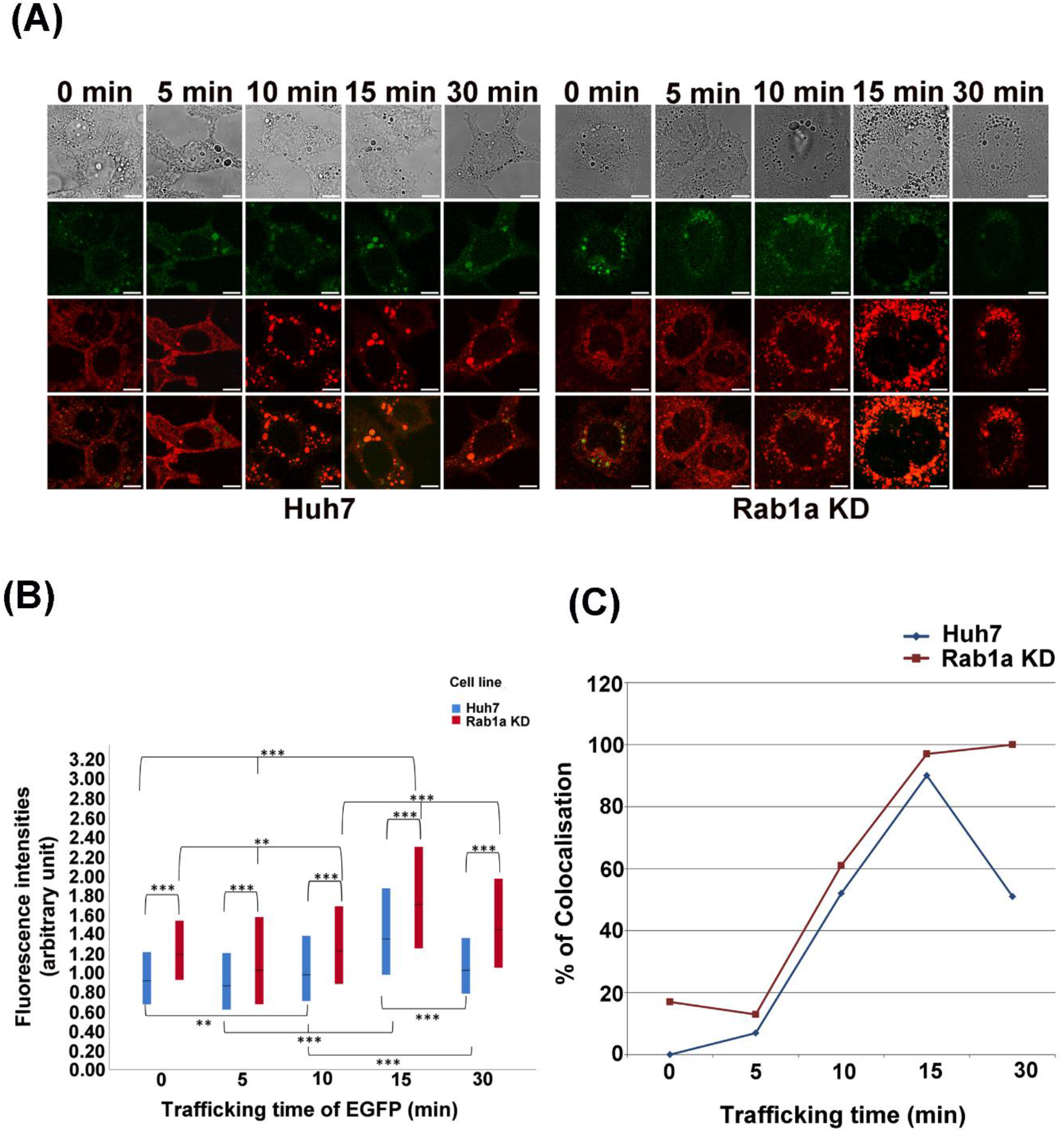
HCV pseudoparticles infection and trafficking in Rab1a KD cells. Representative z-plane confocal images of fixed Rab1a KD Huh7 cells represent time lapse trafficking of HCVpp in Rab1a KD cell line. (A) Representative images of E1E1 GFP (green) trafficking in Huh7 (left) and Rab1a KD (right) cells for indicated time points. Cells were co-stained with FM 4-64 indicated in red. Top panel DIC images, second panel HCVpp trafficking depicted in green. Third panel: FM4-64 staining and bottom panel depict merged images of HCVpp and FM4-64. Scale Bars= 10μm. (B) Integrated density (IntDen) of EGFP from each of the individual infected cells (Huh7 and Rab1aKD) are represented in a box plot profile. Each box represents median, first and third quartile. ** represents significant difference (p < 0.05) and *** represents significant difference (p < 0.001) as tested by ANOVA. For Huh7 (0 min), N= 1748; for 5min, N= 1018; for 10min, N= 768; for 15min, N= 556; for 30min, N= 906 and for Rab1a KD (0min), N=504; for 5min, N=286; for 10min N= 468; for 15min, N= 425; for 30min N= 555.(C) Graphical representation of positive colocalization percentage (%) between GFP and FM 4-64 are observed at indicated time points. For Huh7 cell lines, N (0 min)= 67, N (5 min)= 46, N(10 min)= 33, N(15 min)= 60 and N(30min)= 76; and for Rab1a KD, N(0 min)= 30, N(5min)= 47, N(10 min)= 28, N(15 min)= 74 and N(30 min)= 30. Total number of IntDen measurements for images throughout a z-stack for all replicates of a time point were counted as N. One way ANOVA followed by Bonferonni post hoc analysis was performed for statistical analysis. t-score was determined to check the significance of Pearson’s R (above threshold) values.

**Fig. 7.**
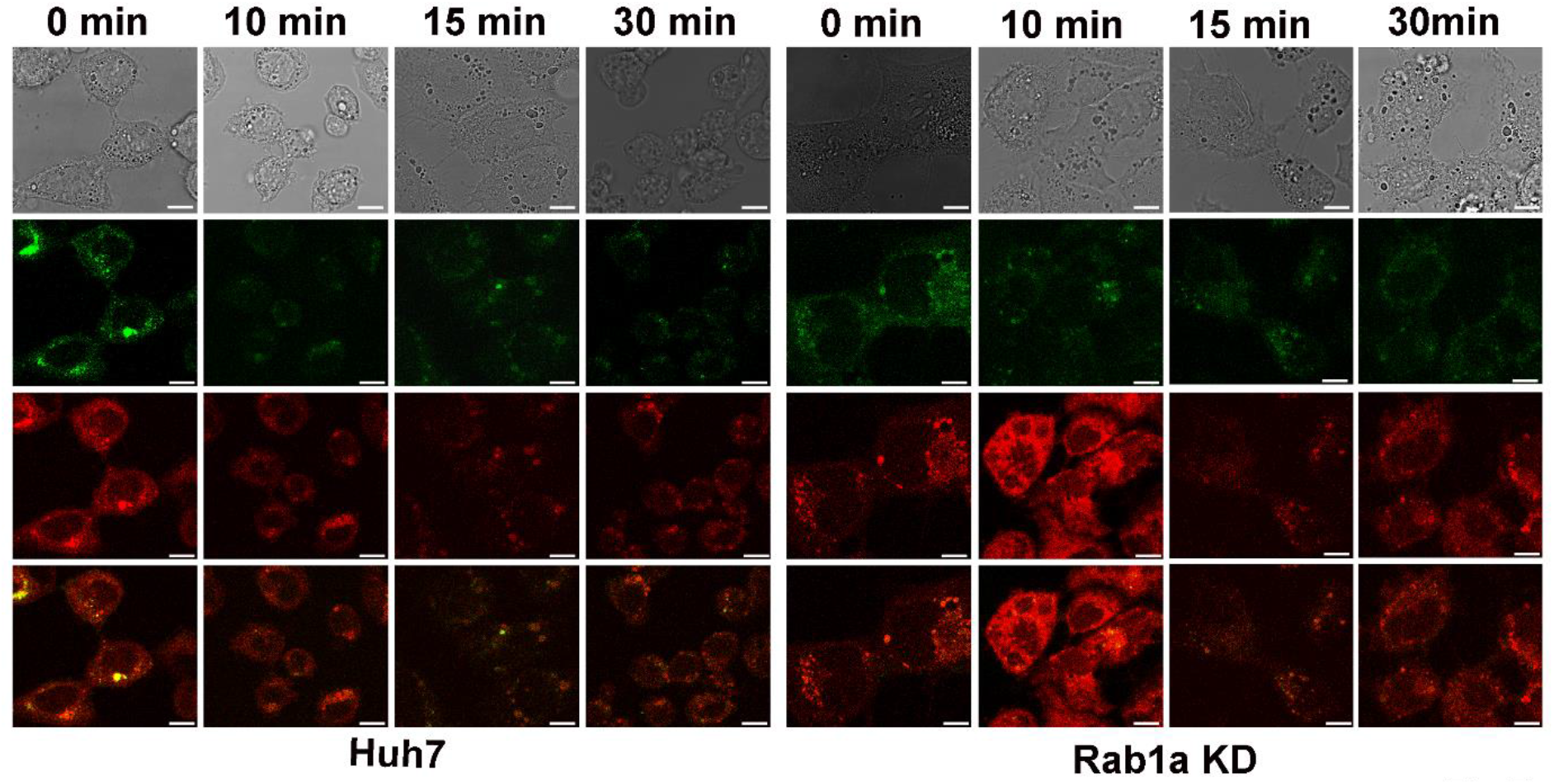
HCVpp fails to reach lysosomes. Representative z-plane confocal images of fixed Huh7 and Rab1a KD cells represent time lapse trafficking of HCVpp in Rab1a KD and Huh7 cell line. HCVpp infected Huh7 and Rab1a KD cells were co-stained with lysosome marker, Lysotracker (depicted in red). Colocalisation of HCVpp (green) and Lysotracker were assayed. Total numbers of cells for each time points are counted as N. For Huh7 cell lines, N (0 min)= 50, N (5 min)= 45, N(10 min)= 45, N(15 min)= 50 and N(30min)= 55; and for Rab1a KD, N(0 min)= 50, N(5min)= 37, N(10 min)= 40, N(15 min)= 55 and N(30 min)= 56.

### 3.4. Rab1a is important for HCVpp trafficking

In order to study the role of cellular proteins in HCV entry using these fluorescent pseudoparticles, the importance of Rab1a in HCV trafficking was studied. Rab1a, apart from its role in ER-Golgi trafficking has also been previously characterized to be important in early endocytic sorting events as well as trafficking of early endosomes by recruiting kinesin motor KifC2 thereby enabling microtubule based movement (Mukhopadhyay et al., 2014). We decided to investigate the role of Rab1a in HCV trafficking by using a Rab1a KD cell line (Mukhopadhyay et al., 2011). Experiments previously described for Huh7 were performed in Rab1a KD cell line and compared to the events occurring in the parental Huh7 cells. Observation of trafficking and measurement of pixel intensity, (Fig. 6) revealed increased pixel intensity at all time points compared to the parental Huh7. This was similar to that observed for trafficking of other ligands in Rab1a KD cells lines and is indicative of delayed trafficking leading to accumulation of ligand. (Fig. 6A). Maximum pixel intensity was observed at 15 minutes and stayed almost constant till 30 min. Furthermore, colocalization with FM 4-64 (Fig. 6B) revealed that while in Huh7 co-localisation with HCVpp reduced between 15-30 minutes, there was no such reduction observed in Rab1aKD cells. As Rab1a is known to slow down endocytic trafficking at early time points, between 5-15 minutes (Mukhopadhyay et al., 2014), we conclude that Rab1a has an important role to play in the endocytic trafficking of HCV. To cross check for an association of E1E2-EGFP with lysosomes in the Rab1a KD cells, we co-stained with Lyso Tracker and did not observe any colocalization (Fig. 7).

## 4. Discussion

In this study we have describe the preparation of fluorescent pseudoparticles which can be used in microscopic assays on fixed cells. These pseudoparticles can possibly be used for live cell imaging as well, which in our preliminary studies proved to be successful (data not shown). To the best of our knowledge this is the first description of fluorescent pseudoparticles which have been used in trafficking assays to determine the very early events of viral entry. In our studies we have described the trafficking of fluorescent HCVpp in Huh7 cells and predict approximate fusion timing. This information helps to understand the timing of viral entry and thereby design methods of intervention. We also describe for the first time a role of the Rab GTPase (Rab1a) in early trafficking events of this virus. Rab 1a has been previously reported to be involved in early endocytic processing of various ligands (Mukhopadhyay et al., 2014), but never known to be important in any viral entry event. This study indicates that Rab1a has a role in HCV trafficking but whether it may be tapped as a potential target to stop HCV entry completely needs to be proved by further experimentation.

This work has far reaching implication beyond the understanding of HCV entry only. These trafficking assays can be used to understand the entry of any pseudotyped virus including SARS CoV-2. Such assays have the potential to be a powerful tool to screen and understand the mechanistic of any preventive of viral entry. Fluorescence labelled lentiviral particles can used to understand viral entry, rate of transmission or most importantly early trafficking details of any virus by simple qPCR, FACS analysis or confocal microscopy.

## Acknowledgements

The authors acknowledges funding from department of Biotechnology-BioCARe Project (BT/Bio-CARe/07/10139/2013-14) awarded to AM and DBT-BUILDER awarded to Presidency University. The authors would also like to acknowledge funding from Presidency University-Faculty Research Professional Development Fund.

## Author Contributions

Chayan Bhattacharjee: Investigation, Formal analysis, Writing-original draft,

Aparna Mukhopadhyay: Conceptualization, Methodology, Writing-Review and Editing, Supervision, Funding acquisition.

## References

Baldick, C.J., Wichroski, M.J., Pendri, A., Walsh, A.W., Fang, J., Mazzucco, C.E., Pokornowski, K.A., Rose, R.E., Eggers, B.J., Hsu, M., Zhai, W., Zhai, G., Gerritz, S.W., Poss, M.A., Meanwell, N.A., Cockett, M.I., Tenney, D.J., 2010. A novel small molecule inhibitor of hepatitis C virus entry. PLoS Pathog. 6. https://doi.org/10.1371/journal.ppat.1001086

Bananis, E., Murray, J.W., Stockert, R.J., Satir, P., Wolkoff, A.W., 2000. Microtubule and motor-dependent endocytic vesicle sorting in vitro. J Cell Biol 151, 179–186.

Bartosch, B., Cosset, F.L., 2009. Studying HCV cell entry with HCV pseudoparticles (HCVpp). Methods Mol. Biol. 510. https://doi.org/10.1007/978-1-59745-394-3_21

Bartosch, B., Dubuisson, J., Cosset, F.-L., 2003. Infectious hepatitis C virus pseudo-particles containing functional E1-E2 envelope protein complexes. J. Exp. Med. 197, 633–42.

Blanchard, E., Belouzard, S., Goueslain, L., Wakita, T., Dubuisson, J., Wychowski, C., Rouille, Y., Rouillé, Y., 2006. Hepatitis C Virus Entry Depends on Clathrin-Mediated Endocytosis. J. Virol. 80, 6964–6972. https://doi.org/10.1128/JVI.00024-06

Diao, J., Pantua, H., Ngu, H., Komuves, L., Diehl, L., Schaefer, G., Kapadia, S.B., 2012. Hepatitis C Virus Induces Epidermal Growth Factor Receptor Activation via CD81 Binding for Viral Internalization and Entry. J. Virol. 86, 10935–10949. https://doi.org/10.1128/JVI.00750-12

HIV-based vectors. Preparation and use - PubMed [WWW Document], n.d. URL https://pubmed.ncbi.nlm.nih.gov/11987783/ (accessed 1.9.21).

Hsu, M., Zhang, J., Flint, M., Logvinoff, C., Cheng-Mayer, C., Rice, C.M., McKeating, J.A., 2003. Hepatitis C virus glycoproteins mediate pH-dependent cell entry of pseudotyped retroviral particles. Proc. Natl. Acad. Sci. U. S. A. https://doi.org/10.1073/pnas.0832180100

Lavanchy, D., 2009. The global burden of hepatitis C, in: Liver International. https://doi.org/10.1111/j.1478-3231.2008.01934.x

Mukhopadhyay, A., Nieves, E., Che, F.-Y.Y., Wang, J., Jin, L., Murray, J.W.W., Gordon, K., Angeletti, R.H.H., Wolkoff, A.W.W., 2011. Proteomic analysis of endocytic vesicles: Rab1a regulates motility of early endocytic vesicles. J Cell Sci 124, 765–775. https://doi.org/10.1242/jcs.079020

Mukhopadhyay, A., Quiroz, J.A.A., Wolkoff, A.W.W., 2014. Rab1a regulates sorting of early endocytic vesicles. Am J Physiol Gastrointest Liver Physiol 306, G412–24. https://doi.org/10.1152/ajpgi.00118.2013

Murray, J.W., Sarkar, S., Wolkoff, A.W., 2008. Single vesicle analysis of endocytic fission on microtubules in vitro. Traffic 9, 833–847.

Saunier, B., Triyatni, M., Ulianich, L., Maruvada, P., Yen, P., Kohn, L.D., 2003. Role of the Asialoglycoprotein Receptor in Binding and Entry of Hepatitis C Virus Structural Proteins in Cultured Human Hepatocytes. J. Virol. 77. https://doi.org/10.1128/jvi.77.1.546-559.2003

Voisset, C., Op de Beeck, A., Horellou, P., Dreux, M., Gustot, T., Duverlie, G., Cosset, F.L., Vu-Dac, N., Dubuisson, J., 2006. High-density lipoproteins reduce the neutralizing effect of hepatitis C virus (HCV)-infected patient antibodies by promoting HCV entry. J. Gen. Virol. 87. https://doi.org/10.1099/vir.0.81932-0

Willingham, M.C., Pastan, I.H., 1982. Transit of epidermal growth factor through coated pits of the Golgi system. J Cell Biol 94, 207–212.

